# Generation and characterization of a novel MHC-II tetramer for tracking and characterization of toxin B-specific CD4^+^ T cell responses

**DOI:** 10.64898/2026.02.18.706639

**Authors:** Jeffrey R. Maslanka, Qianxuan She, Kathleen S. Krauss, Emily N. Konopka, Nile U. Bayard, Jennifer Londregan, Mohamad-Gabriel Alameh, Laurence C. Eisenlohr, Michele A. Kutzler, Joseph P. Zackular, Michael C. Abt

**Author notes:** Corresponding Author: Department of Microbiology, Perelman School of Medicine, University of Pennsylvania, Philadelphia, PA, USA. 303B Johnson Pavilion, 3610 Hamilton Walk. Philadelphia, PA, 19104.

## Abstract

The gastrointestinal pathogen *Clostridioides difficile*, is a major burden for health systems due to high rates of recurrence. *C. difficile* pathogenesis is mediated by two virulence factors, toxin A (TcdA) and Toxin B (TcdB). Antibodies specific for TcdA and TcdB are correlated with protection from symptomatic recurrence, however, the role for CD4^+^ T cells is poorly understood in part due to the lack of tools to study the toxin-specific CD4^+^ T cell response. Our group recently demonstrated the antibody and CD4^+^ T cell response to *C. difficile* toxins is impaired via the glucosyltransferase activity of the toxins; however, tools do not exist to study the protective capacity and the phenotype of toxin-specific CD4^+^ T cells. Therefore, we developed an MHC-II tetramer to identify TcdB-specific CD4^+^ T cells via flow cytometry. Herein, we identified an immunodominant epitope (TcdB_1961-1975_) in the CROPs region of TcdB and optimized an MHC-II tetramer for use in tracking and phenotyping TcdB-specific CD4^+^ T cell responses following multiple different immunization strategies in mice. Utilizing the tetramer, TcdB-specific T follicular helper (Tfh) cells were detected following TcdB-CROPs mRNA-LNP vaccination validating the advantage of the tetramer. Furthermore, using a modular mRNA vector expressing the TcdB_1961_ peptide covalently bound to the beta chain of MHC-II (MHC-IIβ) we were able to generate a robust population of TcdB-specific CD4^+^ T cells. These data outline the generation of new tools for the *C. difficile* field and lay the groundwork for future studies of toxin-specific CD4^+^ T cell responses.

## Introduction

*Clostridioides difficile* infection is particularly challenging to manage due to exceptionally high rates of recurrent infections. Multiple studies have identified antibody titers to *C. difficile* toxins are correlated with lower recurrence rates^1–8^, suggesting that antibodies have the capacity to protect from a symptomatic recurrence, however, a subset of patients fail to generate toxin specific antibodies and are therefore at an elevated risk of recurrence. CD4^+^ T cells and their contribution to establishing protective immunity against *C. difficile* recurrence is not well understood. One study by Cook *et al.* identified a reduction in circulating TcdB-responsive T helper 17 (Th17) cells among recurrent patients compared to new onset infection^9^ suggesting TcdB-specific CD4^+^ T cell responses may support protection from recurrence. Mechanistic studies in the murine *C. difficile* infection model to define the contribution of CD4^+^ T cells to protective immunity have also been limited to phenotypic characterization of polyclonal CD4^+^ T cell populations. CD4^+^ T cells robustly expand during *C. difficile* infection, with significant increases in Th17^10^ cells and regulatory T cells (Tregs) in the large intestine lamina propria^11^. Although CD4^+^ T cells are expanded following infection, they are dispensable for survival from acute *C. difficile* infection^11,12^. Furthermore, whether the antigen specificity of these responding CD4^+^ T cells is to *C. difficile* antigen or bystander commensal bacteria and their role to preventing recurrence is largely unknown. The lack of immunologic tools to identify antigen-specific CD4^+^ T cell responses is a major limiting factor for the study of CD4^+^ T cells in murine models. Recently, our group established an *ex vivo* restimulation assay to identify toxin-responsive CD4^+^ T cells utilizing peptide libraries of TcdA and TcdB to identify cytokine producing, CD4^+^ T cells. No significant TcdB-responsive effector CD4^+^ T cell induction was detected following infection with wild-type *C. difficile-*infected mice compared to uninfected controls. Conversely, infection with glucosyltransferase mutant strains of *C. difficile* yielded IFN-γ and IL-17A producing effector CD4^+^ T cells in the large intestine following TcdB-restimulation^10^. These data demonstrated that the effector CD4^+^ T cell response to toxins is inhibited by the glucosyltransferase activity of the secreted toxins^10^. However, assessing CD4^+^ T cell responses by peptide restimulation is limited to cytokine producing effector CD4^+^ T cells and is thus limited in phenotyping and tracking these toxin-specific CD4^+^ T cells *in vivo*. Therefore, better tools are needed to study toxin-specific CD4^+^ T cell responses to enhance our understanding of their role in natural and vaccine induced immunity against *C. difficile*. Herein, a 15 amino acid sequence in the CROPs domain of TcdB was identified as an immunodominant CD4^+^ T cell epitope. This epitope was used to generate and validate a Major Histocompatibility Complex class II (MHC-II) tetramer for the identification and characterization of TcdB-specific CD4^+^ T cell responses following vaccination. Following immunization with a TcdB mRNA lipid nanoparticle (mRNA-LNP) vaccine, mice generate robust TcdB-specific CD4^+^ T cell responses with a subset of TcdB-specific T follicular helper (Tfh) cells. Moreover, Immunization with a TcdB RBD DNA vaccine elicited TcdB-responsive CD4^+^ T cells that were primarily TcdB_1961_-specific Th1 cells responses indicating immunodominance was not unique to mRNA vaccination. Furthermore, immunization with an mRNA-LNP encoding the TcdB_1961_ peptide covalently bound to MHC-IIβ lead to a robust expansion of TcdB-specific Th1 cells. We report these results for the *C. difficile* immunology community to use this tool to conduct future mechanistic studies on the role of toxin-specific CD4^+^ T cells in host immunity against *C. difficile* infection.

## Results

### TcdB_1918_ and TcdB_1961_ are immunodominant epitopes following vaccination

Previous work in our lab demonstrated that despite predicted immunogenicity, and robust population level Th17 responses, *C. difficile*-infected mice failed to elicit TcdB-responsive CD4^+^ T cells unless the glucosyltransferase activity of the toxin was mutated^10^. However, our analysis was limited to TcdB peptide-responsive CD4^+^ T cells producing cytokine. Therefore, additional tools are required to further characterize the TcdB-specific CD4^+^ T cell response. We sought to develop an MHC-II tetramer to identify immunodominant TcdB-specific CD4^+^ T cells by flow cytometric analysis. Potential immunodominant epitopes in our TcdB peptide library were identified using predicted immunogenicity analysis from the immune epitope database (IEDB)^13,14^ on the TcdB-CROPs peptide library generated previously^10^. Six epitopes were identified that were predicted to be highly immunogenic in the context of I-Ab, the MHC-II molecule expressed in C57BL/6 mice (**Figure 1A**). To validate the computational results, a prime/boost TcdB-CROPs mRNA-LNP vaccination strategy previously found to elicit TcdB-responsive CD4^+^ T cells was utilized^15^. Splenocytes were isolated from naïve and vaccinated C57BL/6 mice and stimulated *ex vivo* with the TcdB-CROPs peptide pool or individual peptides identified by our computational analysis, and cells were assessed by flow cytometry for cytokine production. Following stimulation with TcdB pooled peptide, vaccinated mice generated robust polyfunctional Th1 skewed TcdB-responsive CD4^+^ T cell responses compared to unvaccinated mice producing both IFN-γ and TNF-α (**Figure 1B,C**). Next, the cytokine responses to individual peptides were assessed. Peptide number TcdB-083 (TcdB_1918-1932_ TSDGYKYFAPANTVN) and peptide number TcdB-126 (TcdB_1961-1975_ TDEYIAATGSVIIDG) saw significant cytokine production following individual peptide restimulation (**Figure 1B,C**). Therefore, TcdB_1918_ and TcdB_1961_ were determined to be immunodominant CD4^+^ T cell epitopes among the TcdB protein.

**Figure 1.**
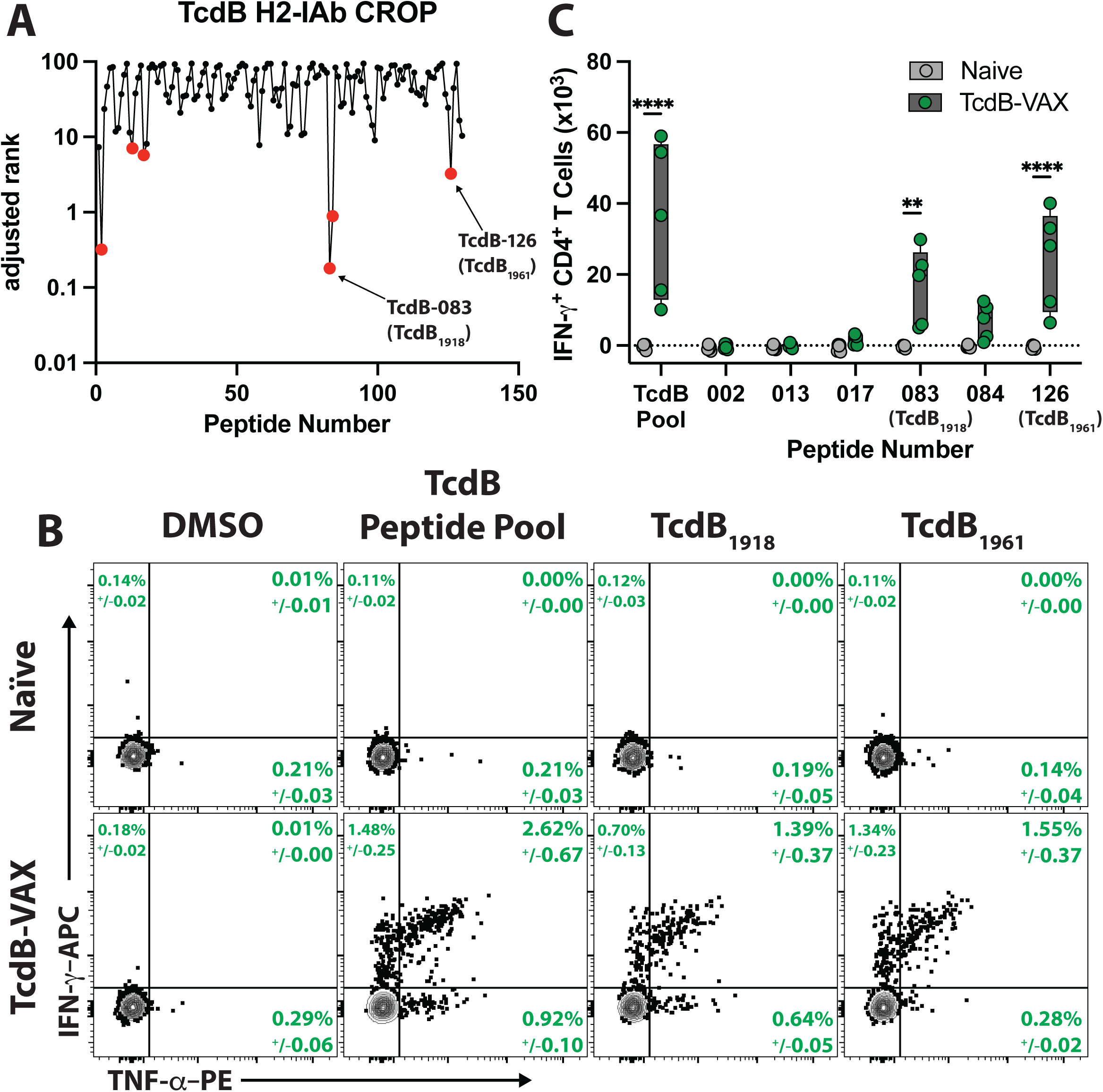
**TcdB_1918_ and TcdB_1961_ are immunodominant epitopes following vaccination.** (A) Amino acid sequences from TcdB peptide library^10^. 15mers were analyzed using the Immune epitope database MHC-II peptide binding prediction for H2-IAb. Data were represented as adjusted rank which is generated by comparing each peptide’s score to a database of 15mers. Lower numbers are predicted to be better binders to H2-IAb. (B) Representative FACS plots showing frequency of TcdB pooled peptide library, individual peptides, or DMSO stimulated CD4^+^ T cells expressing IFN-γ. (C) The total number of IFN-γ ^+^ CD4^+^ T cells responding to TcdB peptide library or individual peptides. To account for the background cytokine production observed, the total number of IFN-γ ^+^ CD4^+^ T cells in the DMSO group was subtracted from the total number of peptide-responsive IFN-γ ^+^ CD4^+^ T cells. Naïve n=5 and TcdB-CROPs mRNA vaccinated n=5 per timepoint. Two-way ANOVA with Tukey’s multiple comparison test. Statistical significance is indicated as follows: ** P < 0.01; **** P < 0.0001. Representative FACS plots were pre-gated based on the following parameters: Singlets, Live, Lymphocytes, CD45^+^, CD3/5^+^, CD4^+^ CD44^+^, CD62L^lo^. Frequency of parental gate +/− standard error of the mean is shown in green. Cells were stimulated with peptide or DMSO for 1 hour before the addition of BFA-M. Cells were incubated for an additional 4 hours before intracellular cytokine staining and analysis by flow cytometry.

### TcdB_1918_ and TcdB_1961_ are conserved and unique to toxigenic *C. difficile*

TcdB has been demonstrated to be hypermutated among *C. difficile* strains with multiple different toxin types expressed across various clades of *C. difficile*^16^. Therefore, identification of a conserved epitope is critical for characterization of immune responses to all *C. difficile* isolates. Following identification of two potential epitopes, TcdB_1918_ and TcdB_1961_ were compared to a database of *C. difficile* isolates^17^. TcdB_1918_ and TcdB_1961_ both had regions within the phylogenic tree of high and low percent identity, however, all strains with low percent identity were subsequently determined to be non-toxigenic (**Figure 2A**). Next, toxigenic and non-toxigenic strains were grouped and compared to a database of gut bacteria to determine if the TcdB_1918_ and TcdB_1961_ epitopes were unique to *C. difficile*. Toxigenic *C. difficile* isolates had both high percent identity and percent coverage, while no gut bacteria had both high percent identity and percent coverage to either TcdB_1918_ or TcdB_1961_ (**Figure 2B,C**). Therefore, TcdB_1918_ and TcdB_1961_ are unique to *C. difficile* and conserved among toxigenic *C. difficile* strains. Both experimental and computational data confirm that these are promising epitopes to generate an MHC-II tetramer to detect CD4^+^ T cells with TCRs that are specific for TcdB.

**Figure 2.**
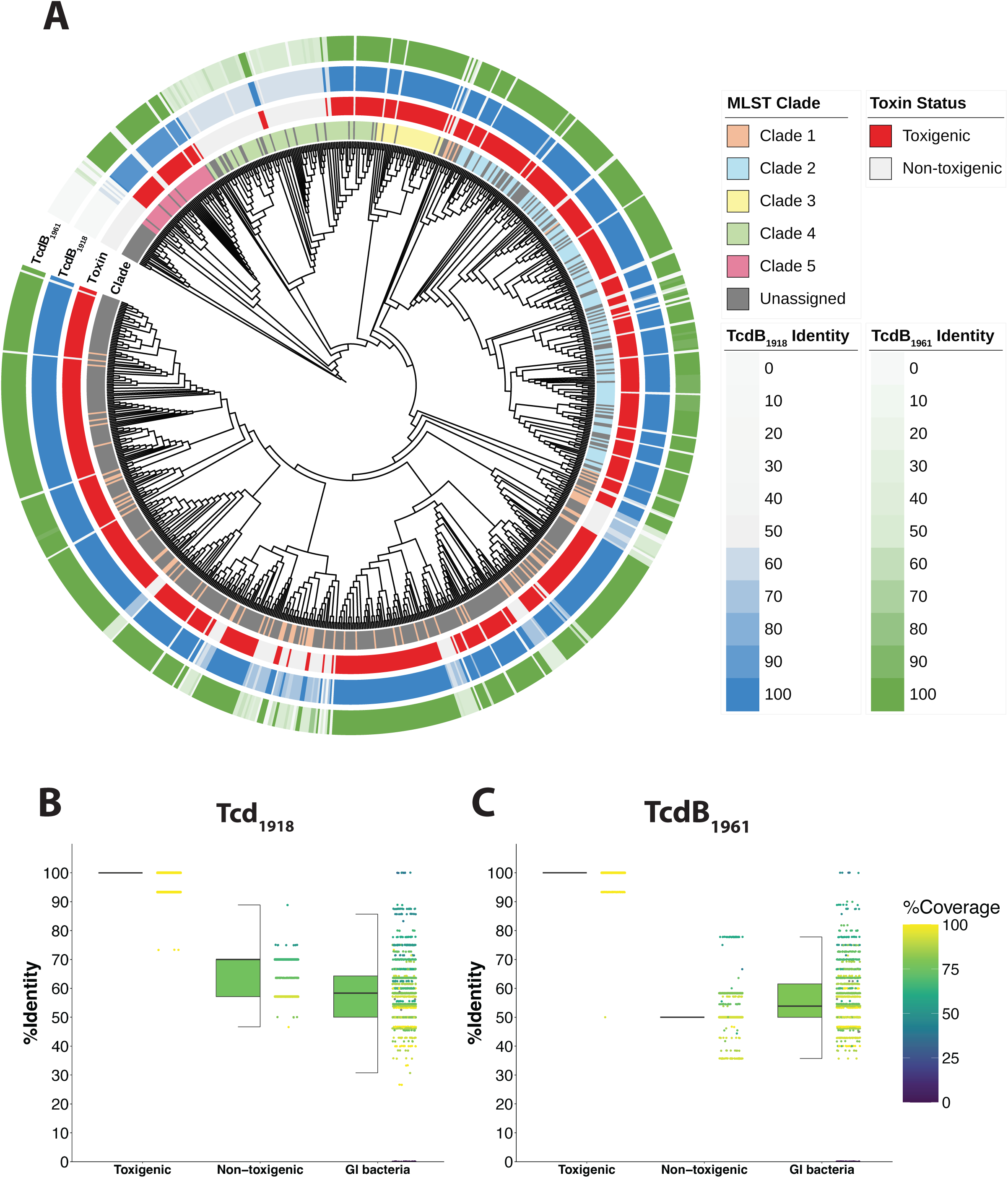
**TcdB_1918_ and TcdB_1961_ are conserved among toxigenic *C. difficile* strains and unique to toxigenic *C. difficile*.** (A) Phylogenetic tree of 923 representative *C. difficile* genomes with four annotation rings. From outermost to innermost: (1) percent amino acid identity to TcdB_1961_ (green gradient, 0–100%); (2) percent amino acid identity to TcdB_1918_ (blue gradient, 0–100%); (3) toxin status, indicating toxigenic (red) or non-toxigenic (light grey) classification, and (4) MLST clade assignment. **(B–C)** Distribution of percent amino acid identity to TcdB_1918_ **(B)** and TcdB_1961_ **(C)** across toxigenic *C. difficile*, non-toxigenic *C. difficile*, and other gastrointestinal bacterial species. Half-boxplots (left) display the median (center line), interquartile range (IQR; box bounds), and 1.5× IQR (whiskers). Individual data points (right) are jittered for visualization, with color indicating percent of coverage (viridis scale, 0–100%). Boxplot fill color represents mean coverage for each group.

### Optimization of the TcdB_1961_ IA-b MHC-II tetramer

TcdB_1918_ and TcdB_1961_ tetramers were both developed through the service of the NIH tetramer core facility, however, the TcdB_1961_ tetramer demonstrated increased signal compared to the TcdB_1918_ tetramer in initial testing (data not shown) and therefore is the focus of the remainder of this report. To determine the optimal incubation temperature, time, and concentration for the TcdB_1961_ tetramer to bind to toxin-specific CD4^+^ T cells, mice were vaccinated and boosted utilizing the same TcdB-CROPs mRNA-LNP vaccine strategy^15^. Splenocytes were incubated in the presence of the TcdB_1961_ tetramer or a negative control CLIP tetramer at 4°C and 37°C. Incubating at 37°C significantly enhanced the frequency and total number of TcdB_1961_ tetramer^+^ CD4^+^ T cells detected in vaccinated mice compared to controls demonstrating that 37°C was required for optimal tetramer staining (**Figure 3A,B**). Next, tetramer concentration was assessed. The TcdB_1961_ tetramer was titrated and incubated at 37°C. Higher concentration of TcdB_1961_ tetramer significantly enhanced the frequency and total number of TcdB_1961_ tetramer^+^ CD4^+^ T cells in vaccinated mice compared to controls demonstrating that high concentration was required for optimal TcdB_1961_ tetramer staining (**Figure 3C,D**). Finally, incubation time was assessed. Splenocytes from naïve and vaccinated mice were incubated with high concentrations of TcdB_1961_ tetramer at 37°C. Two-hour incubation led to an increase in the frequency of TcdB_1961_ tetramer^+^ CD4^+^ T cells in vaccinated mice but no increase in total number was observed indicating that longer incubation times were beneficial but not required for optimal tetramer staining (**Figure 3E,F**). Incubation of control CLIP tetramer at these various conditions revealed a slight increase in control tetramer binding at 37°C and higher concentrations (**Supplemental Figure 1**), however, the increased TcdB_1961_ tetramer staining at these conditions demonstrate these conditions yield the optimal signal for TcdB_1961_ tetramer staining.

**Figure 3.**
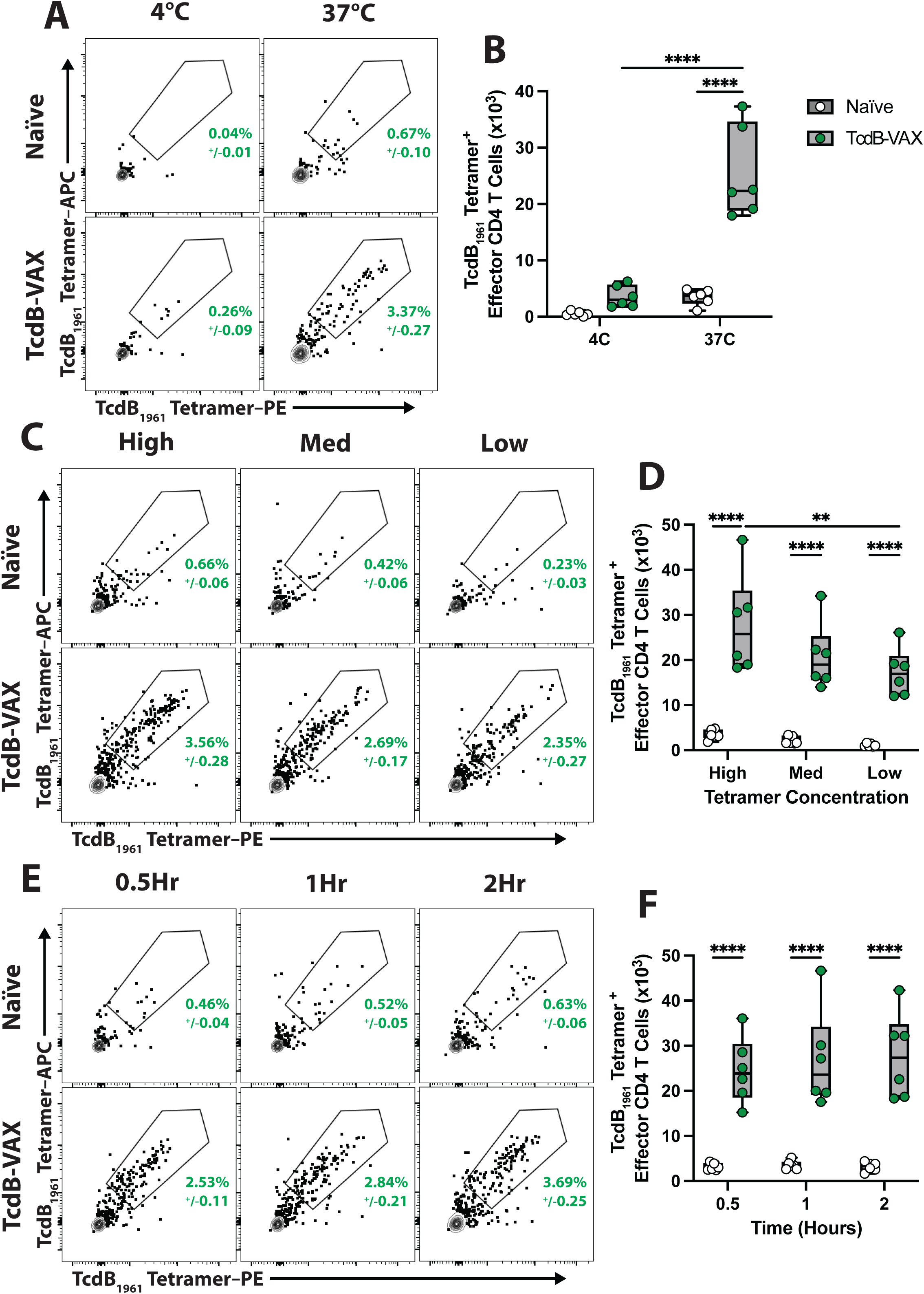
**37**°**C incubation and high concentrations are required for optimal tetramer signal.** (A) Representative FACS plots showing frequency and (B) total number of TcdB_1961_ tetramer binding CD4^+^ T cells from naïve and TcdB-CROPs mRNA-LNP-vaccinated mice at various incubation temperatures. (C) Frequency and (D) total number of TcdB_1961_ tetramer binding CD4^+^ T cells at various tetramer concentrations. High: 18µg/mL Med: 7µg/mL Low: 3µg/mL. (E) Frequency and (F) total number of TcdB_1961_ tetramer binding CD4^+^ T cells at various incubation times. Splenocytes from naïve and TcdB-CROPs mRNA-LNP-vaccinated mice were isolated on day 7 post-boost. Cells were fixed using 2% PFA following flow cytometry staining. Representative data of two experiments Naïve n=6 and TcdB-CROPs mRNA vaccinated n=6 per timepoint. Two-way ANOVA with Tukey’s multiple comparison test. Statistical significance is indicated as follows: ** P < 0.01; **** P < 0.0001. Representative FACS plots were pre-gated based on the following parameters: Singlets, Live, Lymphocytes, CD45^+^, CD3/5^+^, CD4^+^ CD44^+^, CD62L^lo^. Frequency of parental gate +/− standard error of the mean is shown in green.

### Fixation protocol enhances signal to noise ratio

Intranuclear (IN) staining of transcription factors is required to identify CD4^+^ T helper subsets. Therefore, an intranuclear fixation/permeabilization buffer was utilized to prepare cells for transcription factor staining and assess how tetramer staining is affected in naïve and vaccinated mice. We observed a significant reduction in the frequency and total number of TcdB_1961_ tetramer^+^ CD4^+^ T cells in vaccinated mice when the intranuclear fixation/permeabilization buffer was used (**Figure 4A,B**). However, when the total number of TcdB_1961_ tetramer^+^ cells was divided by CLIP control tetramer^+^ cells as an indicator for signal to noise ratio, a significant increase in the signal to noise ratio was observed when the intranuclear staining buffer was used compared to cells that were fixed with paraformaldehyde (**Figure 4C**). Furthermore, a significant reduction in CLIP control tetramer binding in both Naïve and vaccinated mice was observed (**Supplemental Figure 2**). Therefore, intranuclear fixation/permeabilization of the cells leads to a reduction in signal but an enhanced signal to noise ratio increasing the accuracy of the TcdB_1961_ tetramer staining.

**Figure 4.**
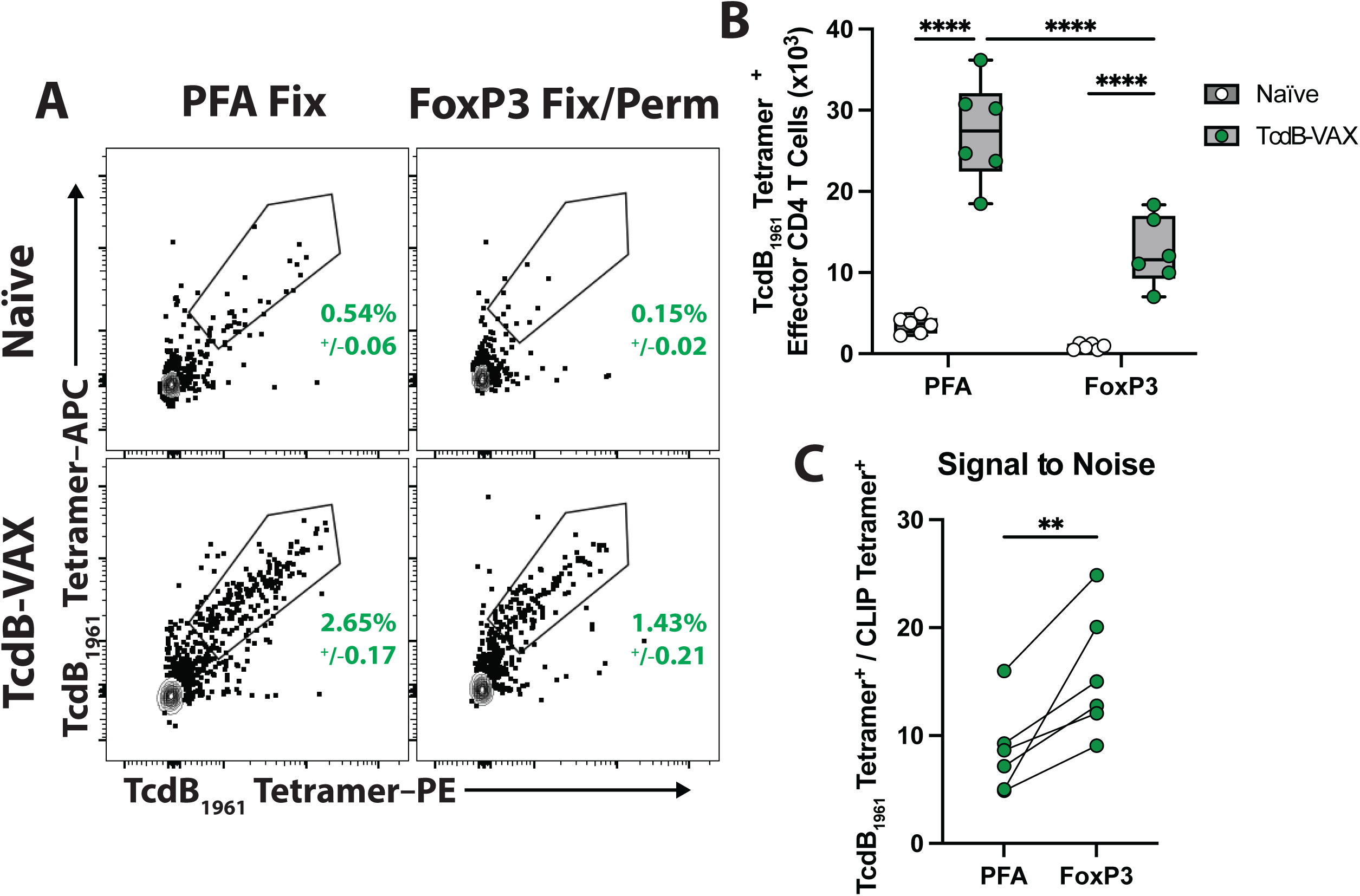
**Intranuclear permeabilization increases signal to noise ratio among tetramer^+^ CD4^+^ T cells.** (A) Representative FACS plots showing frequency and (B) total number of TcdB_1961_ tetramer binding CD4^+^ T cells from naïve and TcdB-CROPs mRNA-LNP-vaccinated mice with different fixation protocols. (C) Signal to noise ratio is the total number of TcdB_1961_ tetramer binding CD4^+^ T cells divided by the total number of CLIP tetramer binding CD4^+^ T cells among TcdB-CROPs mRNA-LNP-vaccinated mice. Splenocytes from naïve and TcdB-CROPs mRNA-LNP-vaccinated mice were isolated on day 9 post-boost. Representative data of two experiments Naïve n=6 and TcdB-CROPs mRNA vaccinated n=6 per timepoint. (B) Two-way ANOVA with Tukey’s multiple comparison test. Statistical significance is indicated as follows: **** P < 0.0001. (C) Paired t test. Statistical significance is indicated as follows: ** P < 0.01. Representative FACS plots were pre-gated based on the following parameters: Singlets, Live, Lymphocytes, CD45^+^, CD3/5^+^, CD4^+^ CD44^+^, CD62L^lo^. Frequency of parental gate +/− standard error of the mean is shown in green.

### TcdB-CROPs mRNA vaccination leads to induction of tetramer^+^ Tfh cells

To verify the utility of the TcdB_1961_ tetramer to identify subsets of CD4^+^ T cells, naïve and vaccinated mice were stained for Tfh markers. At day 9 post-boost, there was no significant difference in frequency or total number of the polyclonal Tfh cell population in the spleen of vaccinated mice (**Figure 5A,B**). However, TcdB_1961_ tetramer staining revealed TcdB_1961_-specific Tfh cells only in mice that received the TcdB-CROPs mRNA-LNP vaccine (**Figure 4C,D**). These data demonstrate that population level changes are not indicative of the antigen-specific response and the TcdB_1961_ tetramer provides a deeper analysis of the CD4^+^ T cell response enabling the investigation of the quality and phenotype of the TcdB-specific adaptive immune response.

**Figure 5.**
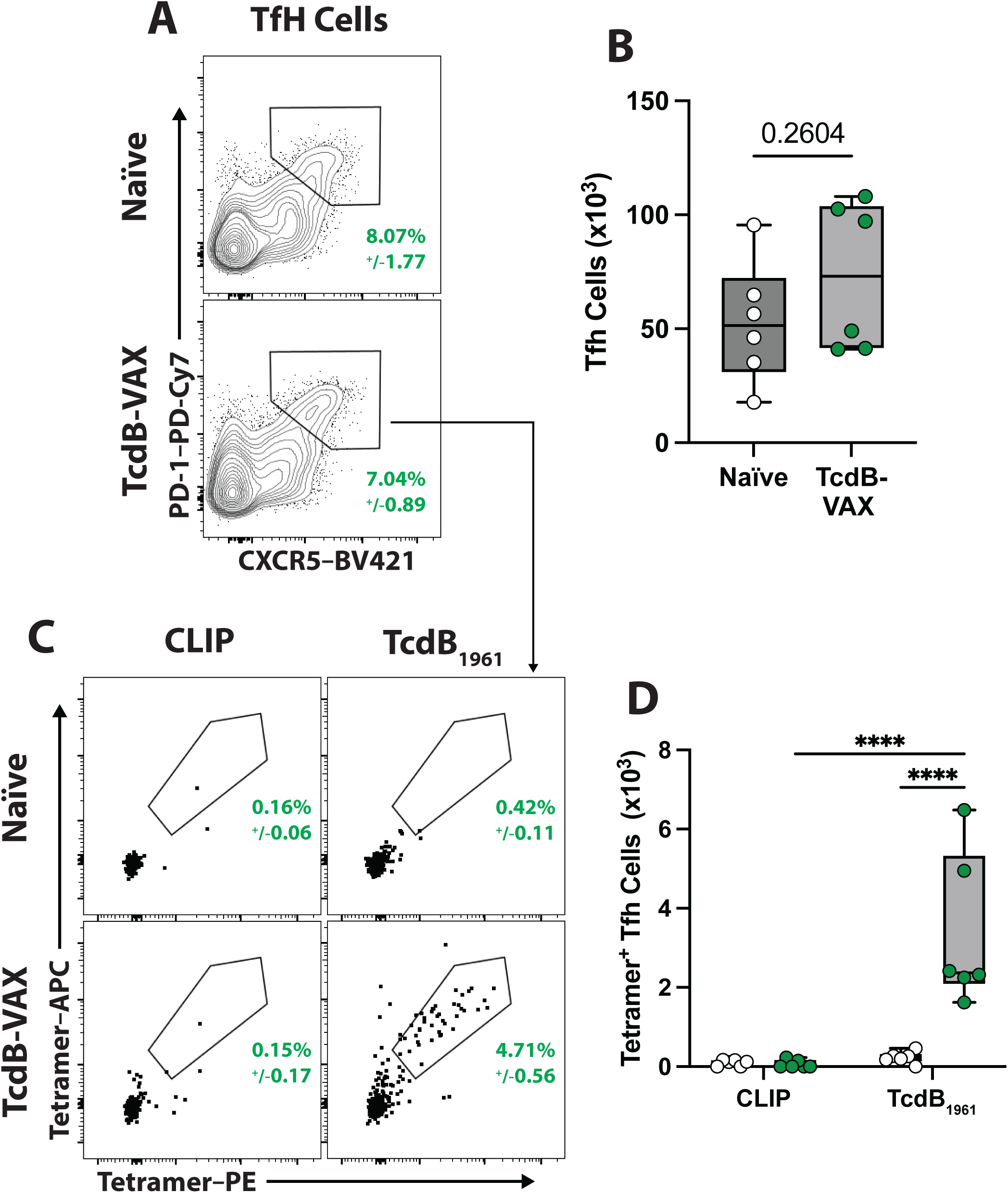
**TcdB-CROPs mRNA-LNP vaccination induces TcdB_1961_-specific Tfh cells.** (A) Representative FACS plots showing frequency and (B) total number of Tfh cells from the spleens of naïve and TcdB-CROPs mRNA-LNP-vaccinated mice. (C) Frequency and (D) total number of TcdB_1961_ tetramer binding Tfh cells. Splenocytes from naïve and TcdB-CROPs mRNA-LNP-vaccinated mice were isolated on day 9 post-boost. Representative data of two experiments Naïve n=6 and TcdB-CROPs mRNA vaccinated n=6 per timepoint. Cells were fixed using 2% PFA or an intranuclear (IN) fixation/permeabilization buffer following surface staining. (B) Unpaired t test. (D) Two-way ANOVA with Tukey’s multiple comparison test. Statistical significance is indicated as follows: **** P < 0.0001. Representative FACS plots were pre-gated based on the following parameters: Singlets, Live, Lymphocytes, CD45^+^, CD3/5^+^, CD4^+^ CD44^+^, CD62L^lo^. Frequency of parental gate +/− standard error of the mean is shown in green.

### TcdB_1961_ tetramer detects CD4^+^ T cell responses across multiple different vaccine platforms

To ensure immunodominance of the TcdB_1961_ epitope was not unique to mRNA-LNP immunization, CD4^+^ T cell responses to a previously reported *C. difficile* toxin receptor binding domain (RBD) DNA vaccine was assessed^18^. Mice were immunized using a prime-boost strategy with a TcdB RBD-encoding plasmid and sacrificed on day 10 post-boost. Splenocytes were stimulated with DMSO, TcdB pooled peptide or TcdB_1961_ peptide to assess CD4^+^ T cell responses following DNA vaccination. CD4^+^ T cells from RBD DNA-vaccinated mice produced IFN-γ in response to the TcdB pooled peptide and the TcdB_1961_ peptide indicating that the TcdB_1961_ epitope remains an immunodominant epitope even in the context of a structurally distinct vaccine modality (**Figure 6A,B**). *Ex* vivo TcdB_1961_ tetramer staining of splenocytes was performed, and significant induction of TcdB_1961_ tetramer^+^ CD4^+^ T cells was observed in DNA-vaccinated mice while CLIP tetramer staining remained identical between groups (**Figure 6C,D**). Together, these data demonstrate that the TcdB_1961_ epitope is reliably targeted across diverse immunization strategies and underscore the utility of the TcdB_1961_ tetramer as a sensitive and specific reagent for quantifying toxin!Zspecific CD4⁺ T cell responses independent of cytokine production.

**Figure 6.**
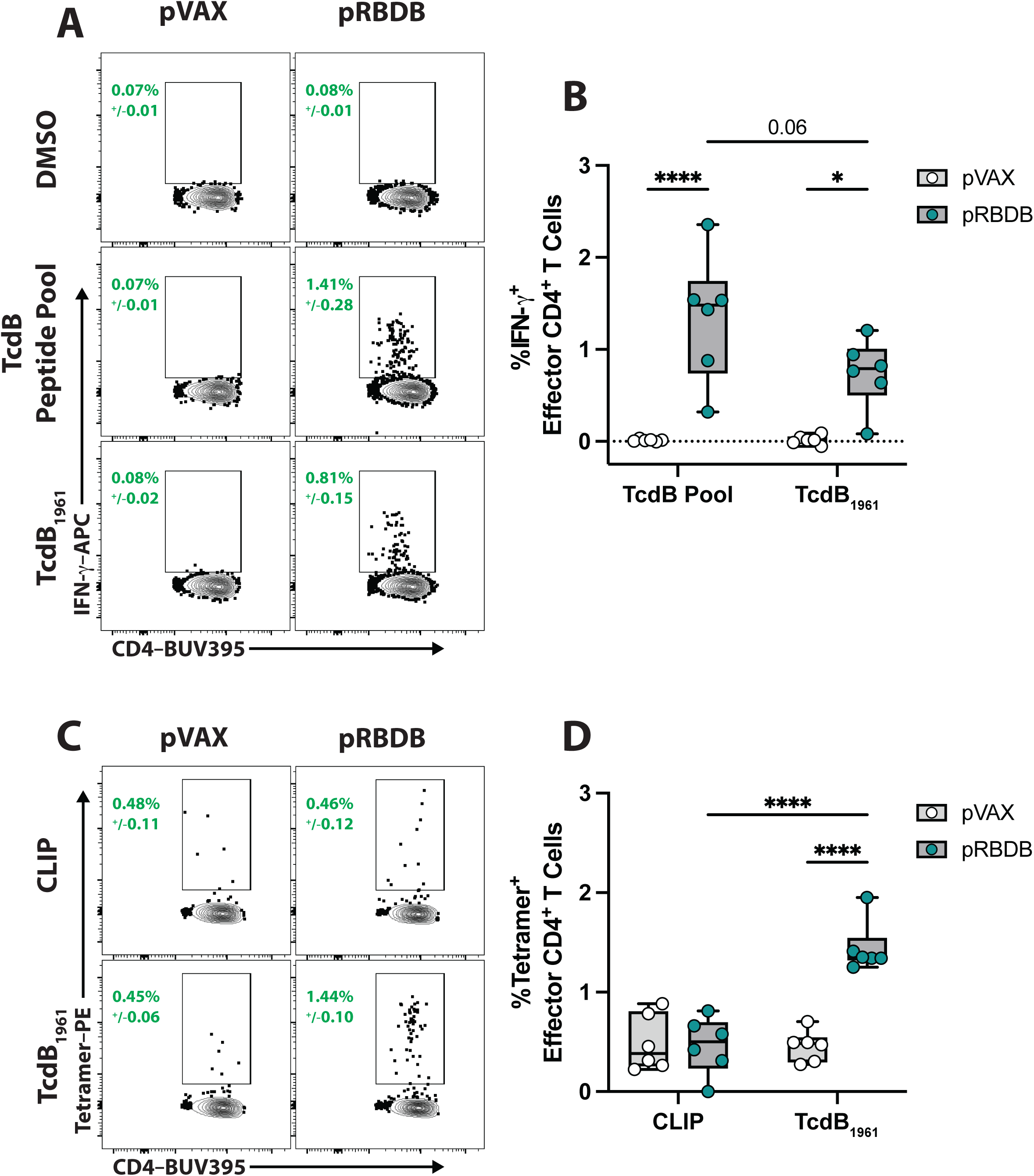
**TcdB_1961_ tetramer works in multiple different vaccination platforms.** (A) Representative FACS plots showing (B) frequency of peptide responsive IFNγ^+^ effector CD4^+^ T cells from the speen of pRBDB DNA vaccinated and pVAX control mice. (C,D) Frequency of TcdB_1961_ tetramer^+^ effector CD4^+^ T cells in vaccinated and control mice. Representative experiment from three replicate experiments. pVAX n=6 and pRBDA pRBDB n=6. Splenocytes from pVAX and pRBDB vaccinated mice were isolated on day 10 post-boost. One-way ANOVA with Tukey’s multiple comparison test. Statistical significance is indicated as follows: **** P < 0.0001. Representative FACS plots were pre-gated based on the following parameters: Singlets, Live, Lymphocytes, CD45^+^, CD3/5^+^, CD4^+^ CD44^+^, CD62L^lo^. Frequency of parental gate +/− standard error of the mean is shown in green.

### TcdB_1961_ IA-b mRNA-LNP immunization induces robust TcdB-specific Th1 responses

To generate, track, and characterize TcdB_1961_-specific CD4^+^ T cell responses, a modular mRNA platform that encodes the TcdB_1961_ peptide covalently linked to the N-terminus of the beta chain of Major Histocompatibility Complex II (MHC-IIβ) was utilized^19^. Mice immunized with the TcdB_1961_/IA-b mRNA-LNP generated a robust TcdB_1961_-specific CD4^+^ T cell response in the spleen (**Figure 7A,B**). Most of the TcdB_1961_-specific CD4^+^ T cells expressed high levels of the transcription factor T-bet indicating they were Th1 cells (**Figure 7B,C**) in agreement with Krauss et al. that reported this modular mRNA platform elicits a strong Th1 response^19^. This proof-of-concept technique demonstrates the ability to selectively generate TcdB_1961_-specific CD4^+^ T cells for use in experimental systems to study the kinetics of toxin-specific CD4^+^ T cells with a defined TCR-specificity.

**Figure 7.**
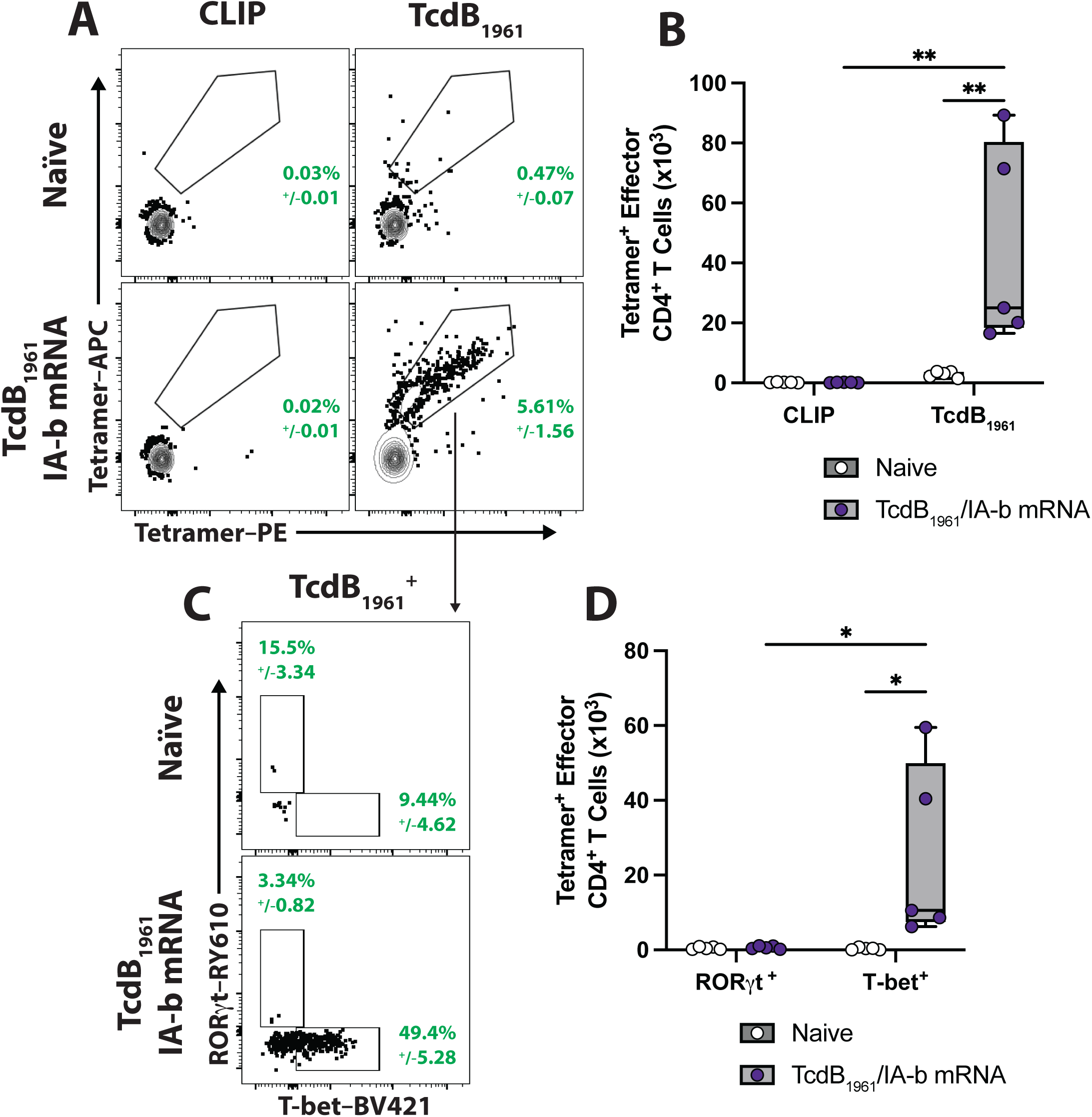
**TcdB_1961_/IA-b mRNA-LNP immunization induces robust TcdB_1961_ specific Th1 cells.** (A) Representative FACS plots showing frequency and (B) total number of TcdB_1961_ tetramer binding CD4^+^ T cells from the spleen of TcdB_1961_/IA-b mRNA-LNP immunized and naïve mice at post-vaccination. (C) Frequency and (D) total number of T-bet^+^ TcdB_1961_ tetramer^+^ CD4^+^ T cells in TcdB_1961_/IA-b mRNA-LNP immunized mice. Naïve n=5 and TcdB_1961_/IA-b mRNA-LNP n=5. Splenocytes from naïve and TcdB_1961_/IA-b mRNA-LNP-immunized mice were isolated on day 4 post-boost. One-way ANOVA with Tukey’s multiple comparison test. Statistical significance is indicated as follows: * P < 0.05; ** P < 0.01. Representative FACS plots were pre-gated based on the following parameters: Singlets, Live, Lymphocytes, CD45^+^, CD3/5^+^, CD4^+^ CD44^+^, CD62L^lo^. Frequency of parental gate +/− standard error of the mean is shown in green.

## Discussion

Immunological research on *C. difficile* pathogenesis has historically been focused on antibody responses and innate immunity with a lack of studies on CD4^+^ T cells, despite the critical role these cells likely play in shaping the neutralizing antibody response and long-term protection from recurrence. A major barrier has been the lack of resources to track and to phenotype toxin-specific CD4^+^ T cell responses. Work from our group has generated peptide libraries for use in *ex vivo* restimulation assays to detect cytokine producing toxin-responsive CD4^+^ T cells^10,15^, however, the field still lacked a way to track toxin-specific CD4^+^ T cell subsets that was not reliant on the cells’ capacity to produce cytokines. Consequently, the antigen specificity, functional heterogeneity, and protective capacity of CD4^+^ T cell responses in *C. difficile* infection have remained largely undefined. Herein, an immunodominant epitope for TcdB was identified that was both unique to *C. difficile* and conserved among toxigenic strains. An MHC-II tetramer was generated to identify CD4^+^ T cells with TCR cognate for this epitope. The tetramer performed strongly across distinct vaccine modalities including an mRNA-LNP platform, a DNA-based toxin RBD vaccine, and an MHC-β linked monospecific mRNA-LNP construct demonstrating its versatility and broad applicability. Notably, a portion of TcdB-specific CD4^+^ T cells generated following TcdB mRNA-LNP vaccination were Tfh cells, a particularly challenging cell type to identify through traditional peptide restimulation methods. Although the total number of Tfh cells did not differ between vaccinated and control mice at this timepoint, tetramer staining revealed that the total number of TcdB_1961_-specific Tfh cells was significantly expanded in vaccinated mice, underscoring that conventional surface staining alone is insufficient to fully and accurately phenotype the cellular immune responses generated following vaccination. These data highlight the utility of the TcdB_1961_ tetramer for characterizing the quality and phenotype of TcdB-specific CD4^+^ T cell responses.

The TcdB_1961_ tetramer described herein opens the field to a wide degree of studies on CD4^+^ T cells in response during both *C. difficile* infection and vaccination. The TcdB_1961_ tetramer enables detailed investigation of TcdB-specific CD4^+^ T cell differentiation pathways, memory formation, Tfh and Treg induction, and potential correlates of protection against recurrence. Because recurrence remains one of the most significant clinical challenges in *C. difficile* infection, tools that allow for direct study of toxin-specific T cell immunity are essential for advancing rational vaccine design and for identifying immune features that predict protective vs pathogenic responses. We propose that the TcdB_1961_ epitope may serve as a model antigen for identification and characterization of toxin specific adaptive immune response for *C. difficile* studies *in vivo*. Furthermore, TcdB_1918_ may serve as an additional epitope for tracking TcdB-specific CD4^+^ T cells.

Experiments and studies on CD4^+^ T cell responses to model antigens typically utilize transgenic mice where every T cell expresses the same TCR specific for the model antigen. We sought to simplify this process through utilization of a recently established modular mRNA system where a peptide of interest is covalently bound within the binding pocket of MHC-IIβ^19^. Herein, TcdB_1961_/IA-b mRNA-LNP was generated and tested. Immunization with the TcdB_1961_/IA-b mRNA-LNP induced robust monospecific CD4^+^ T cell responses to the TcdB_1961_ epitope, generating a pool of TcdB_1961_-specific CD4^+^ T cells without the need for TCR-transgenic mice. This system may serve as a model for generation of TcdB-specific CD4^+^ T cells for phenotypic and functional studies on toxin specific CD4^+^ T cell responses *in vivo* during *C. difficile* infection.

The focus of this report was the TcdB-specific CD4^+^ T cell response. No tetramer for TcdA was produced in the current study though such a resource would be equally valuable for the field. Preliminary observations suggest there is a broader CD4^+^ T cell epitope landscape for TcdA, complicating identification of a representative immunodominant epitope (data not shown). Future work generating a tetramer to detect TcdA-specific CD4^+^ T cells will require careful consideration that any single epitope selected is representative of the total TcdA-specific CD4^+^ T cell response. Nonetheless, the frameworks established here including epitope discovery, tetramer optimization, and use of TcdB_1961_/IA-b mRNA-LNP tools provide a roadmap for generating additional tools for other epitopes within *C. difficile* as well as other pathogens.

Taken together, these findings address a major unmet need in the *C. difficile* field by establishing robust tools for directly tracking toxin-specific CD4^+^ T cell responses. The TcdB_1961_ tetramer and monospecific IA-b mRNA-LNP system expand the immunological toolkit available to the field and will enable mechanistic studies of CD4^+^ T cell function and their role in recurrence. These tools will be critical for turning correlative studies into mechanistic insights into how CD4^+^ T cells influence disease recurrence through integration with high-dimensional immunoprobing. Defining protective CD4^+^ T cell responses will be critical for designing next-generation vaccines that elicit durable protective immunity to *C. difficile*.

## Methods

### Mice

C57BL/6 mice were bred and maintained at the University of Pennsylvania or Drexel University College of Medicine under specific pathogen-free conditions. Mice were maintained in autoclaved cages with autoclaved food and water ad libitum. Sex and age-matched controls were used in all experiments according to institutional guidelines for animal care. All animal procedures were approved by the Institutional Animal Care and Use Committee of the University of Pennsylvania or Drexel College of Medicine prior to initiation of studies.

### Comparative Genomic and Phylogenetic Analysis

#### Selection of Representative *C. difficile* Genomes

A total of 15,673 publicly available *C. difficile* genome assemblies were obtained from the WhatsGNU *C. difficile* database^17^. Genome quality was assessed using CheckM v1.2.2^20^, retaining only assemblies with ≥95% completeness and ≤5% contamination. Genomes were stratified by multilocus sequence typing (MLST) into Clades 1–5 and an unassigned category^21^. To capture genomic diversity while minimizing redundancy, representative genomes were selected using the dRep v3.4.5 genome dereplication module^22^, applying primary and secondary average nucleotide identity (ANI) thresholds of 99% and 99.9%, respectively. This approach yielded 923 representative genomes (**Supplemental Table 1**).

#### Classification of Toxigenic and Non-Toxigenic *C. difficile* Genomes

Toxin gene presence was determined through comparative genomic analysis using the pathogenicity locus (PaLoc) of *C. difficile* strain 630^23^ as a reference. Coding sequences for tcdA, tcdB, tcdC, tcdE, and tcdR were extracted, and the PaLoc boundaries were defined by the flanking genes cdu1 and cdd1. These sequences were used to construct a nucleotide BLAST database^24^. Analysis was performed using BLASTn v2.14.0+^24^, with hits retained if they met the arbitrary thresholds of ≥80% coverage and ≥80% identity. Genomes were classified as non-toxigenic if they exhibited in the 115 bp conserved region and lacked toxin genes or the PaLoc region, and vice versa (raw data was included in **Supplementary Table 2**).

#### Comparative Genomic Analysis of TcdB_1918_ and TcdB_1961_

Amino acid sequences for TcdB_1918_ and TcdB_1961_ were used to construct a protein BLAST database^24^. Comparative analyses were performed using BLASTx v2.14.0+ against both the 923 representative *C. difficile* genomes and 1,316 RefSeq complete genomes representing various gastrointestinal bacterial species from NCBI. For each query, the hits with the highest bit-score were retained, and percent coverage and sequence identity were recorded. **Supplementary Table 3** contains complete BLASTx results along with species classification, representative genome assignment, and NCBI accession numbers for each of the 1,316 bacterial species analyzed.

#### Phylogenetic Analysis of Representative *C. difficile* Genomes

The 923 representative genomes were annotated using Bakta v1.9.2 with Database v5.1^25^. The resulting gff3 files were utilized for core-genome alignment, which was carried out using panaroo 1.5.0^26^. This alignment was then used to infer the maximum likelihood phylogeny in IQ-TREE version 2.3.0^27^, employing the general time-reversible (GTR) substitution model^28^ and accounting for among-site rate heterogeneity using the Gamma distribution and four rate categories^29^. The phylogeny was visualized using iTOL^30^.

### TcdB-CROPs mRNA-LNP vaccine

The TcdB-CROPs mRNA vaccine was generated as previously described^15^. Mice were immunized i.m. with 1μg of mRNA and boosted 21 days later. Mice were euthanized on days 7-10 post-boost and spleens were harvested for analysis.

### DNA plasmid preparation, immunizations, and tissue harvest

Plasmid design, immunizations, and tissue harvest were performed at Drexel University College of Medicine. Plasmids encoding toxin B RBD (residues 1851-2366, pRBDB) sequenced from C. difficile strain VPI10463 were constructed (pVax1 backbone, GeneArt) and optimized as previously described^18^. Optimization of RBD sequences included replacement of asparagine (N-linked glycosylation sites) with glutamine. Young C57BL/6 mice (6-8 weeks old) from Charles River were vaccinated twice over 28 days with DNA plasmids encoding RBDB, or empty pVax plasmid as a control. For vaccinations, mice were injected intramuscularly in the left tibialis anterior muscle with 5µg pRBDB or pVax. The injection site was immediately electroporated using a CELLECTRA electroporator set to two pulses at a constant 0.2Amp current (Inovio Pharmaceuticals). Mice were euthanized 10 days following the second vaccination and spleens were harvested and processed for subsequent T cell analyses. All animals were housed in a temperature-controlled, light-cycled facility at the Drexel University animal care facility (accredited by the Association for Assessment and Accreditation of Laboratory Animal Care). Animal work was performed according to protocols approved by our Institutional Animal Care and Use Committee. Briefly, spleens were individually placed into an automated stomacher for 60 seconds on high and strained through a 70μM strainer to obtain single-cell suspensions. All cell suspensions were counted, and cell viability was determined using a Countess Automated Cell Counter (Invitrogen, Life Technologies). Suspensions were then centrifuged at 500g and resuspended at 10M cells/mL for flow cytometry.

### TcdB_1961_/IA-b mRNA-LNP

The transcript used is a variant of the modular transcript described in Krauss et al,^19^ which allows the expression of an MCHIIβ chain with an attached target epitope. Briefly, the transcript consists of a sequence of regions which encode the following: the native MHCIIβ signal peptide, the targeted epitope, a flexible linker peptide, and the desired MHCIIβ allele. These regions are flanked with untranslated regions, including an encoded poly-A tail. The targeted epitope sequence was derived using the Benchling reverse translation feature to derive a codon-optimized sequence encoding the TcdB epitope. This sequence was cloned into a template plasmid and used for the RNA-LNP production pipeline detailed in Krauss et al., which consists of linearization and purification of the encoding plasmid for use as a template in the T7 Megascript in vitro transcription kit with modifications (the addition of the TriLink CleanCap^®^ reagent and substitution of N1-Methylpseudouridine-Triphosphate for the included uridine triphosphate). The RNA product was then purified with cellulose to reduce dsRNA contaminants^31^. Concentration and purity of the product was assessed by Nanodrop 2000 and gel electrophoresis. The RNA transcript was then encapsulated into lipid nanoparticles (LNPs) as described in Hogan et al^32^. Briefly, citrate-buffered mRNA was rapidly mixed via pipette with a lipid solution of ionizable cationic lipid SM-102 (BroadPharm BP-25499), cholesterol (Avanti 700100P), 1,2-distearoyl-sn-glycero-3-phosphocholine (‘DSPC’, Avanti 850365P) and 1,2-dimyristoyl-rac-glycero-3-methoxypolyethylene glycol-2000 (‘DMG-PEG-2000’, Avanti 880151P) in a 50:38.5:10:1.5 molar ratio. The encapsulated mRNA-LNPs were then dialyzed in PBS overnight. The mRNA-LNPs were cryopreserved by adding sucrose to a final volume of 10%, aliquoted, and stored at −80°C until ready for use, when they were injected i.p. into mice at a dose of 2μg at d0 and d6. Mice were euthanized on day 10 and spleens were harvested for analysis.

### Isolation of immune cells from spleen

Single cell suspensions were obtained from spleen by mechanical separation. Spleens were dissected, placed on a 100µm cell strainer, and crushed with a 3mL syringe to dissociate the tissue. Spleen cells were then ACK treated to lyse red blood cells. Cells were centrifuged at 500g and resuspended in complete tissue culture media (RMPI supplemented with 10% FBS, 1% penicillin/streptomycin, 50 µg/mL gentamicin, 10 mM HEPES, 0.5 mM β-mercaptoethanol, 1mM Sodium Pyruvate, 20 µg/mL L-glutamine) for subsequent analysis.

### Cell stimulation, tetramer labeling, and flow cytometry

To detect cytokine production from CD4^+^ T cells, cells in a single cell suspension were cultured in a flat bottom non-tissue culture treated 96 well plate (Corning) in complete medium. Cells were cultured for 1 hour at 37°C in the presence of TcdA peptide pools, TcdB peptide pools (2.5µg/mL/peptide), with DMSO as a negative control, or PMA (0.05 µg/mL [Sigma]) and Ionomycin (0.5 µg/mL [Sigma]) as a positive control. After 1 hour, Brefeldin-A and Monensin (eBioscience) were added and incubated for an additional 4 hours to allow for cytokine accumulation. Following incubation (or directly from single-cell suspension for surface staining), cells were transferred to 96 well round bottom plates for cell staining for flow cytometry using a standard protocol. For tetramer labeling, single cell suspensions were incubated with tetramer (18 µg/mL) in a 96 well round bottom plate for 2 hours at 37°C before washing and cell staining for flow cytometry using a standard protocol. Briefly, cells were washed in PBS, stained with Zombie UV (BioLegend) at room temperature for 10 minutes, and washed in FACS buffer (PBS, 1% BSA, 50µg/mL Sodium Azide) before blocking at 4°C with anti-CD16/32 antibody (BD Biosciences) and rat IgG (Sigma) for 20 minutes. Next, antibodies against surface antigens were added and stained for 30 minutes at 4°C. Surface antibodies included CD4−BUV395 (BD Biosciences, RM4-5), CD62L−BUV805 (BD Biosciences, MEL-14), CD45−BV605 (BioLegend, 30-F11), CD19−BV650 (BioLegend, 6D5), CD3−PerCP-Cy5.5 (eBioscience, 1452C11), CD5−PerCP-Cy5.5 (eBioscience, 53-7.3), CD44−AF700 (BioLegend, IM7), PD-1−PE-Cy7 (eBioscience, J43). For Tfh staining, cells were incubated with a biotinylated CXCR5 antibody (eBioscience, SPRCL5) for 45 minutes at 4°C and washed before the addition of fluorophore conjugated streptavidin (Streptavidin−BV421 [BioLegend]) with all other surface antibodies. Following surface staining, cells were fixed for 30 minutes at 4°C using 2% paraformaldehyde solution for surface staining only, intracellular cytokine fixation buffer (eBioscience) for intracellular staining, or Foxp3 Transcription Factor Fixation/Permeabilization buffer (eBioscience) for intranuclear staining. Intracellular and intranuclear antibodies were diluted in 1x permeabilization buffer (eBioscience) and stained intracellular antigens for 30 minutes at 4°C. Intracellular antibodies included IFN-γ−APC (eBioscience, XMG1.2) and TNF-α−PE (MP6-XT22), while intranuclear antibodies included FoxP3−FITC (eBioscience, FJK-16s). Cells were washed and then analyzed on a BD Symphony A3 flow cytometer. Flow cytometry data analysis was performed using FlowJo version 10.10. Flow cytometry data for this manuscript were generated in the Penn Cytomics and Cell Sorting Shared Resource Laboratory at the University of Pennsylvania (RRID:SCR_022376). Penn Cytomics is partially supported by the Abramson Cancer Center NCI Grant (P30 016520).

## Data analysis

Data was analyzed using Microsoft Excel version 16.100.2 and transferred into GraphPad Prism version 10.6.0 for data visualization and statistical analysis. One-way ANOVA, two-way ANOVA, or mixed-effects models were utilized depending on the independent variables present and are indicated in figure legends. Statistical significance is indicated as follows: * P < 0.05; ** P < 0.01; *** P < 0.001; **** P < 0.0001. Figures were generated in Adobe Illustrator version 27.7.

## Author Contributions

J.R.M. and M.C.A. contributed to experimental design. J.R.M., K.S.K., E.N.K., N.U.B., and J.L. performed the experiments. J.R.M. and Q.S. Performed computation analysis. M.G.A., L.C.E., M.A.K., and J.P.Z. assisted with reagents, and protocols. J.R.M. and Q.S. analyzed experimental data and generated figures. J.R.M. and M.C.A. wrote the manuscript. All authors have read, discussed, and edited the manuscript and approved the submitted version.

## Acknowledgments

The authors thank members of the Abt Lab for their critical review of the manuscript and thoughtful discussions of the results. We thank the NIH Tetramer Core Facility (NIH Contract 75N93020D00005 and RRID:SCR_026557) for providing the TcdB_1961_ tetramer and the CLIP control tetramer utilized within this manuscript.

## Data availability statement

All data generated or analyzed during this study are publicly available from the corresponding author on upon publication.

## Supplemental Figure Legends

**Supplemental Figure 1. 37**°**C incubation and high concentrations increase background CLIP tetramer staining.** (A) Frequency and (B) total number of CLIP tetramer binding CD4^+^ T cells from naïve and TcdB-CROPs mRNA-LNP vaccinated mice at various incubation temperatures. (C) Frequency and (D) total number of TcdB_1961_ tetramer binding CD4^+^ T cells at various tetramer concentrations. High: 18µg/mL Med: 7µg/mL Low: 3µg/mL. (E) Frequency and (F) total number of CLIP tetramer binding CD4^+^ T cells at various incubation times. Splenocytes from naïve and TcdB-CROPs mRNA-LNP-vaccinated mice were isolated on day 7 post-boost. Cells were fixed using 2% PFA following flow cytometry staining. Representative data of two experiments Naïve n=6 and TcdB-CROPs mRNA vaccinated n=6 per timepoint. Two-way ANOVA with Tukey’s multiple comparison test. Statistical significance is indicated as follows: ** P < 0.01; **** P < 0.0001. Representative FACS plots were pre-gated based on the following parameters: Singlets, Live, Lymphocytes, CD45^+^, CD3/5^+^, CD4^+^ CD44^+^, CD62L^lo^. Frequency of parental gate +/− standard error of the mean is shown in green.

**Supplemental Figure 2. Intranuclear permeabilization decreases CLIP tetramer^+^ binding among CD4^+^ T cells.** (A) Frequency and (B) total number of CLIP tetramer binding CD4^+^ T cells from naïve and TcdB-CROPs mRNA-LNP-vaccinated mice with different fixation protocols. Splenocytes from naïve and TcdB-CROPs mRNA-LNP-vaccinated mice were isolated on day 9 post-boost. Cells were fixed using 2% PFA or an intranuclear (IN) fixation/permeabilization buffer following surface staining. Representative data of two experiments Naïve n=6 and TcdB-CROPs mRNA vaccinated n=6 per timepoint. (B) Two-way ANOVA with Tukey’s multiple comparison test. Statistical significance is indicated as follows: **** P < 0.0001. Representative FACS plots were pre-gated based on the following parameters: Singlets, Live, Lymphocytes, CD45^+^, CD3/5^+^, CD4^+^ CD44^+^, CD62L^lo^. Frequency of parental gate +/− standard error of the mean is shown in green.

**Supplemental Figure 3. Gating strategy.** Representative FACS plots showing gating strategy for effector CD4^+^ T cells. Data shown are spleen cells were isolated from a naïve mouse.

