## Supplementary figures and images for "Generation and characterization of a novel MHC-II tetramer for tracking and characterization of toxin B-specific CD4^+^ T cell responses"

### Supplemental Figures

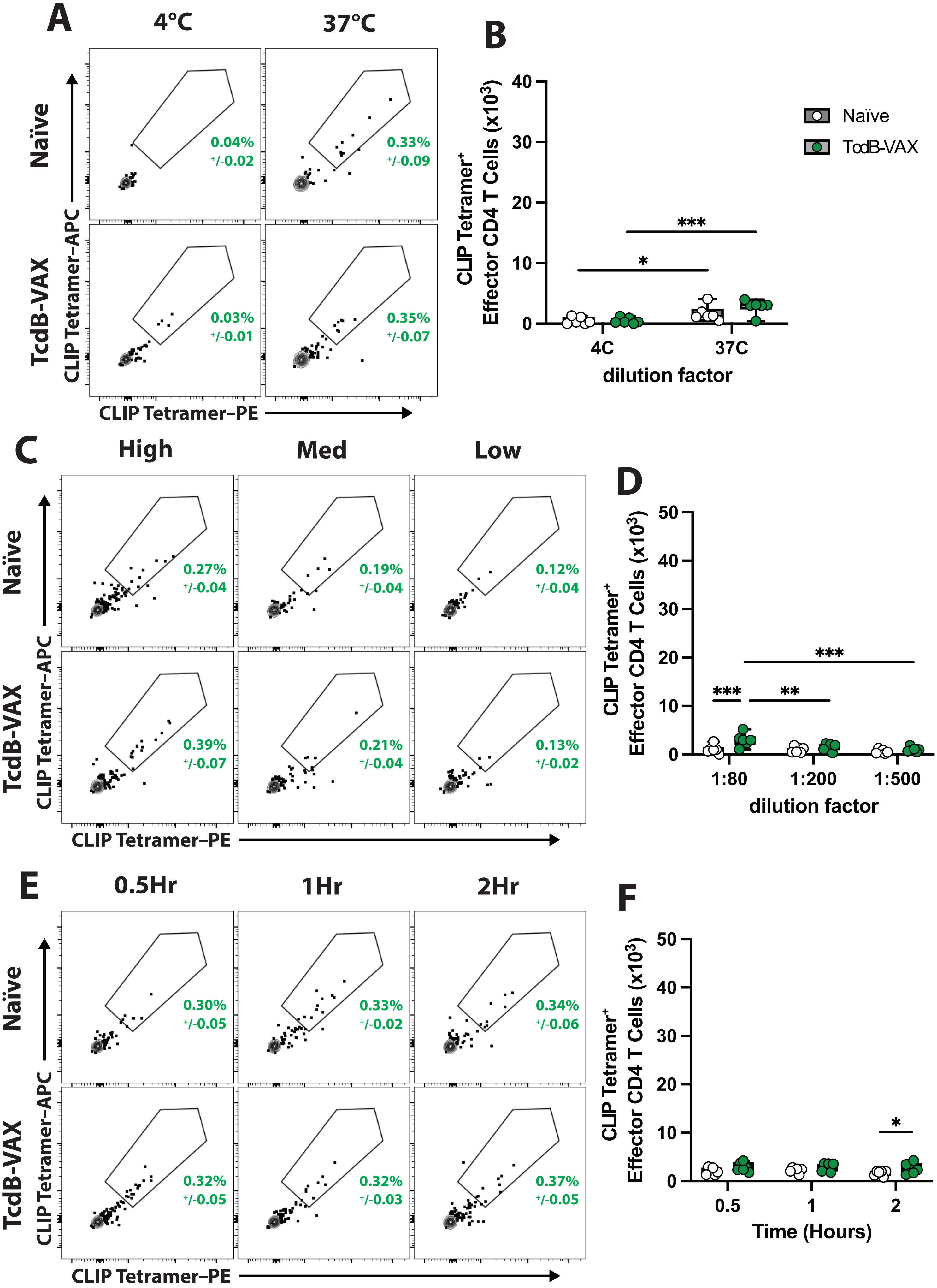

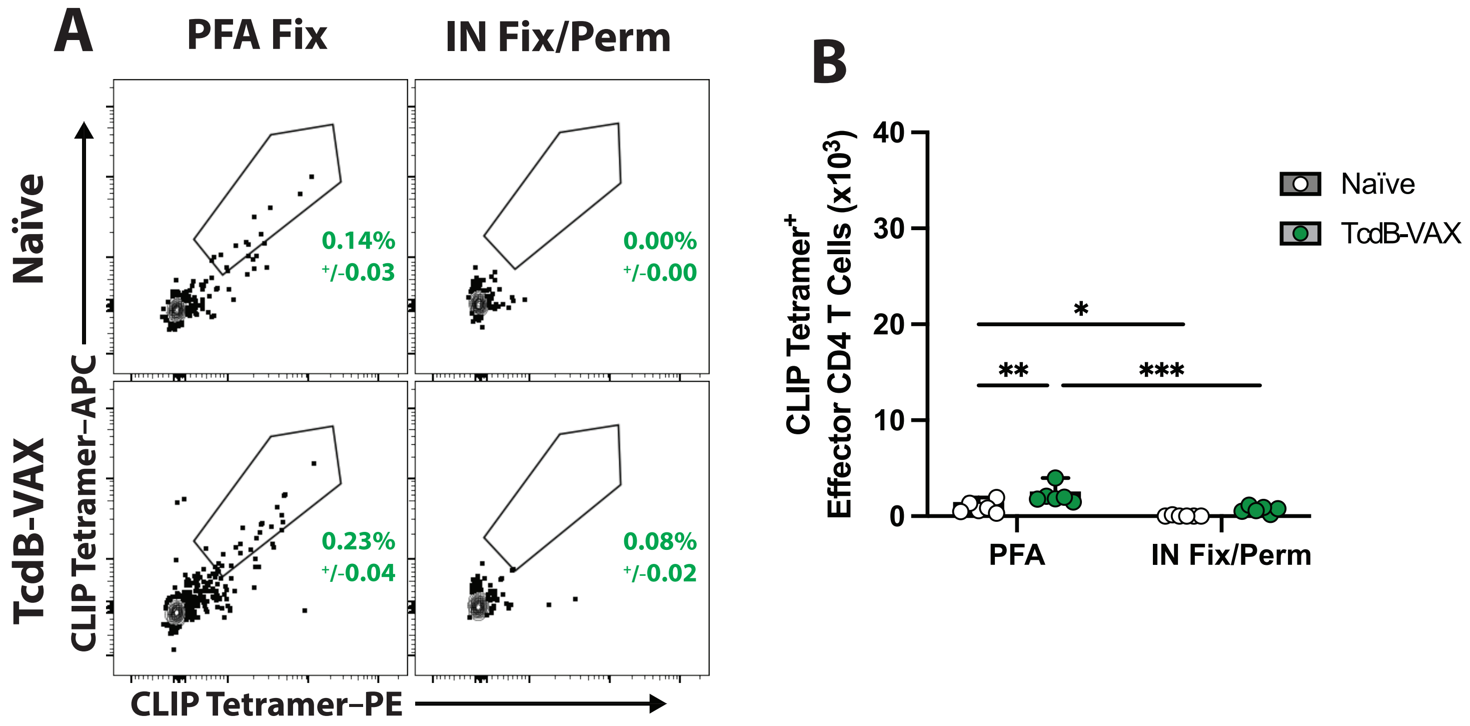

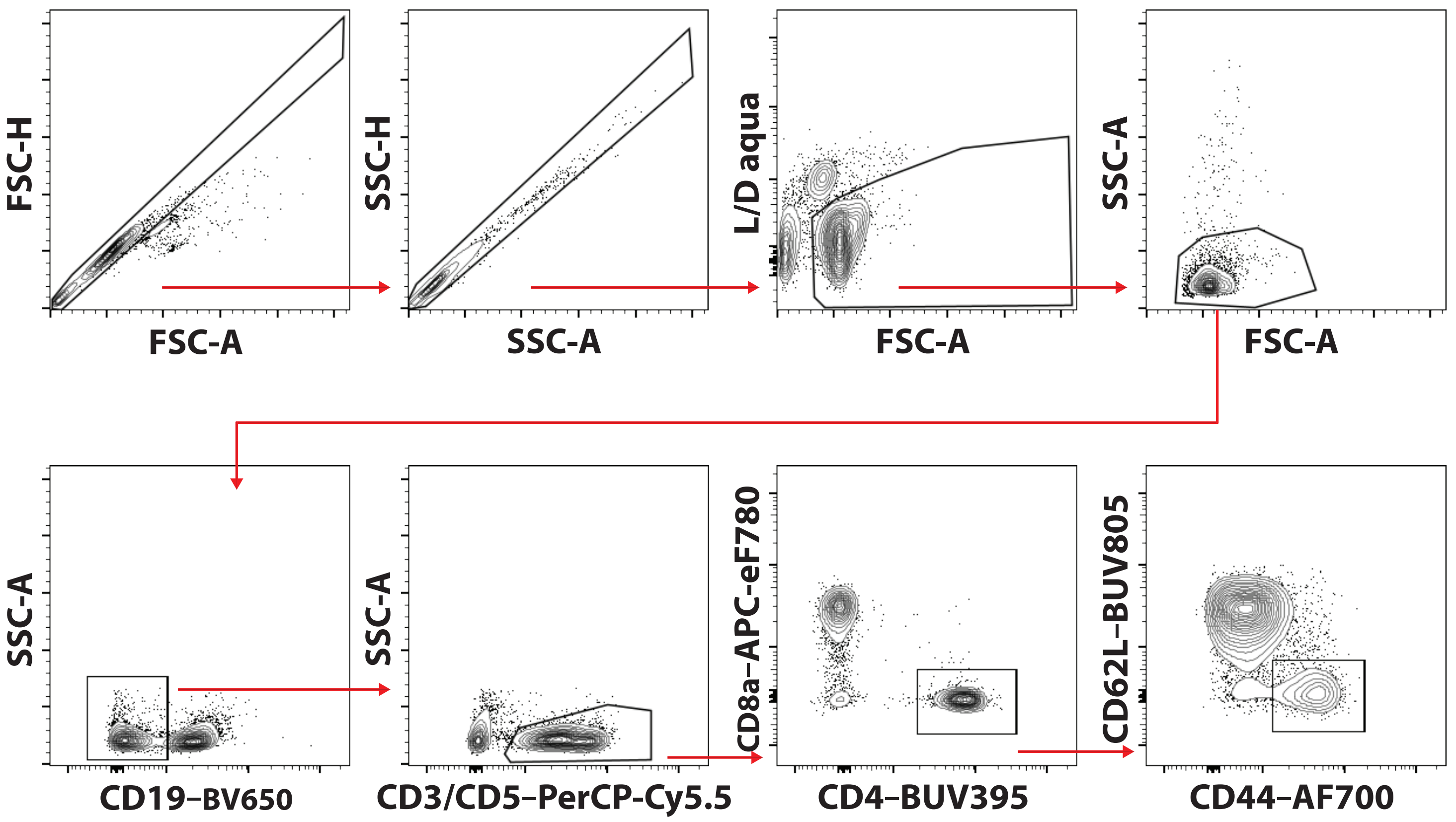
